# Subthalamic neural oscillations reveal no effect of implicit versus explicit facial emotional processing

**DOI:** 10.1101/2020.10.21.348755

**Authors:** Joan Duprez, Thibaut Dondaine, Jean-François Houvenaghel, Julien Modolo, Claire Haegelen, Gabriel Robert, Bruno Millet, Dominique Drapier, Julie Péron, Didier Grandjean, Sophie Drapier, Marc Vérin, Paul Sauleau

## Abstract

In addition to the subthalamic nucleus’ (STN) role in motor control, STN deep brain stimulation (DBS) for Parkinson’s disease (PD) has also uncovered its involvement in cognitive and limbic processing. STN neural oscillations analyzed through local field potential (LFP) recordings have been shown to contribute to emotional (mostly in the alpha band [8-12 Hz]) and cognitive processing (theta [4-7 Hz] and beta [13-30 Hz] bands). In this study, we aimed at testing the hypothesis that STN oscillatory activity is involved in explicit and implicit processing of emotions. To achieve this objective, we used a task that presented patients with fearful emotional facial expressions and asked them to identify the emotion (explicit task) or gender associated with the face (implicit task). We evaluated emotion and task effects on STN neural oscillations power and intertrial phase consistency. Our results revealed that accuracy was lower in the implicit task. Increased STN delta power and decreased alpha and beta power were observed after stimulus presentation. However, there was no influence of emotional facial expression, i.e. neutral versus fear, nor task demands. Intertrial phase consistency in the delta and theta band increased after stimulus onset, in the same time-period as delta power increased. However, similarly to oscillatory power, no changes related to emotional fear expression or task demand were found.

These findings suggest that STN oscillatory activity is not specifically involved in explicit and/or implicit processing of emotions, and that power and phase synchronization changes might be more related to overall task-execution mechanisms. These conjectures remain to be confirmed.

**Highlights:**

– STN LFPs were recorded during an emotional/gender recognition task in PD patients.
– STN delta power increased, and alpha and beta power decreased after stimulus onset.
– Power changes were not influenced by emotional fearful expression or task demands.
– Delta/theta intertrial phase consistency increased after stimulus onset.
– Intertrial phase consistency was not affected by emotional valence or task demands.
– The observed STN activity was likely related to general task-execution mechanisms.

## 1. Introduction

Deep brain stimulation (DBS) of the subthalamic nucleus (STN) is a well-recognized neurosurgical treatment for advanced Parkinson’s disease (PD). Aside from its beneficial, therapeutic effects on motor symptoms, STN-DBS has also been associated with detrimental cognitive (Elgebaly et al., 2018; Odekerken et al., 2015) and emotional effects (Péron et al., 2012). For instance, some studies have pointed out a deficit of facial emotion recognition following STN-DBS (e.g., Biseul et al., 2005; Drapier et al., 2008; Dujardin et al., 2004; Schroeder et al., 2004). The STN is divided in motor, associative and limbic territories (Plantinga et al., 2018), and because of its structural connections and functional properties, it maintains links with numerous cerebral regions involved in the processing of cognitive and emotional information.

The different roles of the STN have also been notably supported by the investigation of its electrical activity through local field potential (LFP) recordings, which allow a direct measurement of the nucleus activity. These measurements are achieved with the DBS electrodes to perform recordings during the time period between electrode implantation and subcutaneous stimulator implantation. LFP recordings during task completion allowed the investigation of STN electrical activity, notably in the time-frequency domain, which describes the dynamics of neural oscillations involved in different situations and frequencies. For instance, abnormal motor activity has been associated with increased STN beta (13-30 Hz) oscillations power (Kühn et al., 2005; Little and Brown, 2014). Regarding cognitive functioning, situations of greater cognitive control are associated with a systematic increase in STN theta (4-7 Hz) power (Duprez et al., 2019; Zavala et al., 2015), and top-down attentional mobilization for relevant information was linked with increased beta power (Engel and Fries, 2010; Zavala et al., 2015).

Regarding emotional processing, a decrease in alpha (8-12 Hz) power followed passive viewing of emotional pictures (Brücke et al., 2007; Kühn et al., 2005) and was further shown to be modulated by depressive symptoms (Huebl et al., 2011). The STN appears to have a specific role in the processing of valence of emotional stimuli, since pleasant pictures (as compared to unpleasant pictures) are associated with a decrease in alpha power (Brücke et al., 2007; Huebl et al., 2014). The role of alpha activity has recently been further confirmed by Mandali et al (2021), who showed that alpha stimulation of the STN was associated with emotional processing. They showed that a 10 Hz stimulation of the right STN was followed by an increase of positive rating after negative stimuli presentation, which was not the case for 130 Hz stimulation. STN involvement in arousal induced by emotional pictures is more nuanced, with reports of both arousal-specific activity (Sieger et al., 2015) and its absence (Brücke et al., 2007). All these studies have pointed out the involvement of STN neural oscillations, especially in the alpha band, in visual emotional processing. However, none of them have specifically focused on STN activity during emotional face recognition, since facial stimuli are often merged with many emotional stimuli of various natures.

Emotion and cognition, although relating to different concepts, are not easily separable in terms of behavior and neural substrates, and are in constant interaction (Dolan, 2002; Ochsner and Phelps, 2007; Pessoa, 2008). Human faces carry relevant biological and social signals which should be processed rapidly to adapt behavior. Considering the functional overlapping territories of the STN and its role in integrating motor, cognitive, and emotional information; this nucleus could arguably participate in the interaction between emotion and cognition. One method to assess this interaction relies on contrasting the activity of this structure between explicit and implicit emotional tasks. In explicit tasks, attention is allocated to emotional information. On the contrary, emotional implicit tasks are defined as emotional processing when attention is oriented towards another characteristic of the stimulus (e.g., gender)., *i.e.*, when there is no conscious awareness of emotional processing (Carretié, 2014; Cohen et al., 2016; De Houwer et al., 2009; D’Hondt et al., 2016; Moors and De Houwer, 2006; Shahane et al., 2019). While the aforementioned studies have shown that the STN is involved in explicit emotional recognition, there is still no electrophysiological study focusing on facial emotion recognition and considering the involvement of the STN in implicit emotional processing.

In order to evaluate the STN’s role in emotional face recognition, and more specifically in implicit emotional processing, we recorded LFPs from the STN during a task based on emotional faces recognition, in which we modulated attention orientation using implicit (gender task) and explicit (emotional discrimination task) instructions. Although this task involved interaction between emotion and cognition in the sense that attention relates to broad cognition, the following results cannot be interpreted as if the task fully segregated emotion from cognition. We expected to record an activity differentially modulated by emotional faces compared to neutral faces (neutral *versus* fearful emotional expression) and attention orientation (implicit *versus* explicit task). Since both emotional- and cognitive-related activity were expected, several frequencies were hypothesized to be modulated during the task, especially in the theta, alpha and beta bands. More precisely, we expected emotional contrast to be associated with changes in alpha activity, and cognitive contrast with theta and beta activity. Furthermore, we aimed at investigating the synchronization of STN neural oscillations across trials, and whether this synchronization was modulated by task factors.

## 2. Methods

### 2.1. Patients and surgery

Sixteen patients with idiopathic PD undergoing bilateral STN DBS at Rennes University Hospital took part in this study (N=16, 9 women). This patient group is the same as in (Duprez et al., 2019), but focusing on a different cognitive task. Patients underwent STN DBS following disabling motor symptoms uncontrolled by optimal pharmacological therapy, and were selected following standard criteria (Welter et al., 2002). Neuropsychological testing (described in details in Péron et al., 2017) indicated that patients were free from major attentional or executive disorders, and did not have deficits in face recognition as estimated with the Benton facial recognition test (Benton et al., 1994). Two patients were excluded from further behavioral and LFP analyses as a result of artifacted data. Thus, the final number of analyzed patients with usable datasets was 14. These patients’ clinical characteristics are presented in Table 1.

**Table 1:** Patients’ preoperative clinical characteristics (average ± standard deviation).

|  |  |
| --- | --- |
| N | 14 |
| Age (years) | 52.6 ± 7.1 |
| Education level (years) | 12.5 ± 3.4 |
| Sex (W/M) | 7/7 |
| Disease lateralization (R/L) | 8/6 |
| Disease duration (years) | 9.4 ± 2.4 |
| UPDRS-III ON dopa | 10.6 ± 7.5 |
| UPDRS-III OFF dopa | 32.3 ± 5.4 |
| S&E ON dopa (%) | 84.6 ± 13.9 |
| S&E OFF dopa (%) | 69.2 ± 20.2 |
| H&Y ON dopa | 1 ± 1 |
| H&Y OFF dopa | 2.3 ± 0.7 |
| LEDD at the time of recording (mg/day) | 800.8 ± 378.4 |
| MDRS (/144) | 140.8 ± 3.1 |
| Benton face recognition test | 44.7 ± 3.4 |

**Abbreviations:** W: Women; M: Men; R: right; L: left; UPDRS: Unified Parkinson's Disease Rating Scale; S&E: Schwab & England; H&Y: Hoehn & Yahr; LEDD: levodopa-equivalent daily dose; MDRS: Mattis Dementia Rating Scale.

Surgery was performed under local anesthesia, using MRI determination of the target, intraoperative microrecordings and assessment of the clinical effects of stimulation. Electrodes with four platinum-iridium cylindrical surfaces (1.27 mm in diameter, 1.5 mm in length, contact-to-contact separation of 0.5 mm; 3389; Medtronic, Minneapolis, MN, USA) were implanted bilaterally in all patients. Implantation was performed so that the lowest contact was positioned at the level of the STN ventral border to use the above contacts to stimulate the sensorimotor part of the nucleus, or at the interface between the subthalamic region and the zona incerta. The first contact (contact 0) was the most ventral, and the fourth (contact 3) was the most dorsal. Correct electrode position was verified post-operatively using a 3D CT brain scan.

This study was conducted in accordance with the declaration of Helsinki, and was approved by the Rennes University Hospital ethics committee (approval number IDRCB: 2011-A00392-39). All patients gave their informed written consent after a thorough description of the study.

### 2.2. Experimental task

The test consisted of two tasks of face recognition always presented in the same order, which differed by the focus of attention (gender vs. emotion). The first task was an implicit emotional faces recognition task (with a gender categorization), and the second task was an explicit emotional faces recognition task (with an emotional labeling). For the implicit emotional task, although faces could express fear or a neutral expression (which was not specified to patients), patients were instructed to answer the question “Did the face represent a man or a woman?” by pressing with their left/right index a left/right button figuring the indication “Man” or “Woman”. For the explicit emotional task, patients were instructed to answer the question “Did the face display fear or a neutral expression?” by pressing with their left/right index a left/right button figuring the indication “Fear” or “Neutral”. We used a set of 80 colored pictures adapted from the Karolinska Directed Emotional Faces database with 20 men and 20 women (Lundqvist et al., 1998) displaying two emotional expressions (fearful or neutral). Non-facial features (hair, neck and clothes) were removed by applying a 250 x 350 pixels elliptical mask. Since these parameters were not controlled in the original version, the luminance and spatial frequencies of each stimulus were normalized using Matlab (Delplanque et al., 2007).

Subjects were seated comfortably in a quiet room in front of a 23’’ computer screen. Stimuli were displayed using E-prime 2.0 Professional edition software (Psychology Software Tools Inc, Sharpsburg, PA, USA). Each trial began with a fixation cross displayed in the center of the screen for a pseudorandomized duration of 700-1500 ms. A face (either expressing a fearful or neutral expression) was then displayed for 250 ms, followed by a black screen for 750 ms. Then, a black screen displaying “Response?” prompted the patient to respond and remained on the screen until a response was given. Finally, the response was followed by a 2000 ms black screen before the next trial began.

Patients were instructed to respond as accurately as possible, without any instruction related to the speed of the response. Each task (implicit/explicit) began by a training session of 20 trials (with different face stimuli than those used in the experimental session). Each task included 80 trials, with an equal proportion of neutral expressions (figured by 20 man faces and 20 women faces) and fearful expressions (figured by 20 man faces and 20 women faces). The same faces were used for the two tasks. All participants were naïve about the purpose of the paradigm that aimed to contrast STN activity when focusing explicitly or implicitly on emotion recognition.

### 2.3. Recordings

All patients were tested two days after surgery, before the implantation of the subcutaneous stimulator. All tests and recordings were performed while patients were under their usual medication. STN LFPs were recorded by a g.BSamp® (g.tec Medical Engineering, Schiedlberg, Austria) biosignal amplifier connected to a PowerLab® 16/35 (ADInstruments, Dunedin, New Zealand) system. For each DBS electrode, activity was recorded bipolarly from all of two adjacent contacts, leading to 3 possible bipolar derivations per electrode: 0-1, 1-2, and 2-3. Signals were amplified and sampled at 1000 Hz, and monitored online using the Labchart® software (ADInstruments). Triggers corresponding to the task stimuli were sent by the Eprime software to the recording device through a parallel port.

### 2.4. Behavioral analyses

All behavioral data analyses were performed using R (version 4.0.2; R Core Team, 2020) implemented with the {lme4} package (Bates et al., 2015). Since no instructions relative to the speed of the response were provided to patients, we only focused behavioral analyses on responses accuracy. The effect of emotional expression (fear, neutral) and task (implicit, explicit) was studied using a mixed-model logistic regression using the *glmer*{lme4} function. Model selection involved testing several models with emotion and task as fixed effects, and adding random intercepts and random slopes for the task effect and emotion effect. We used the Akaike Information Criterion (AIC) for model selection. This criterion informs about the quality of the model, with the lowest value indicating better quality. Thus, the model with the lowest AIC was selected. In this case, the following model was selected that used a *logit link* function and the *bobyqa* optimizer:

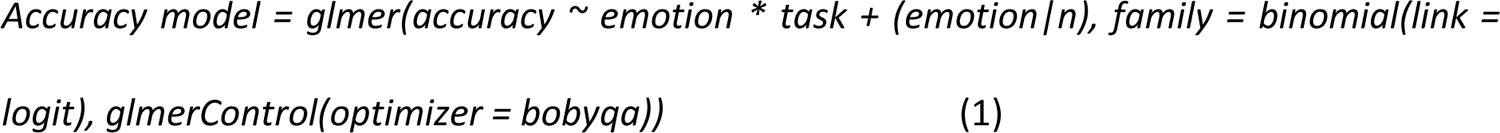

The significance of fixed effects was evaluated by the *Anova*{car} function that provides p-values based on Wald chi-squared tests. We used mixed-models due to their advantages such as the ability to take all data into account, without the need to average at the subject-level, even in the case of unbalanced data; and also due to their ability to model random effects such as subjects (see Gueorguieva and Krystal, 2004). Marginal and conditional R² were obtained from the {MuMin} (Barton, 2009) package. The significance statistical threshold was set at p = 0.05.

### 2.5. LFP signal analyses

All preprocessing steps were carried out using the EEGLAB (Delorme and Makeig, 2004) toolbox for Matlab (The Mathworks, USA). All subsequent time-frequency analyses were performed with custom Matlab code (available at https://github.com/jduprez) based on published equations (Cohen, 2014).

#### 2.5.1. Preprocessing

LFP signals were high-pass filtered offline at 0.5 Hz and epoched from −1 to 2 s surrounding the stimulus (face) display. All epochs were visually inspected and manually removed in the case of excessive noise or artifacts. After preprocessing, 36(±3) trials per experimental condition were available on average. Two patients were excluded from further analyses as a result of excessive noise, resulting in 28 usable STN datasets (two per patient).

Contact pair selection was the result of a two-step procedure. First, anatomical location and CT scan showed that, for all patients, the most distal contact pair (0-1) was located in the ventral part of the STN (1.5 ± 4.2 mm lateral to the anterior-posterior commissure (AC-PC) line, −3.1 ± 1.7 mm posterior to the middle of the AC-PC line, and −5.4 ± 1.7 mm below this point) which has been preferentially associated with cognitive processing (see Supplemental Figure S1 for an example of electrode positioning). Second, a trial-by-trial breakdown (random selection of 25 trials, see Supplemental figure S2) for the three pairs of contacts across all experimental conditions showed that activity was greater for the most distal pair of contacts (0-1), and decreased over the above pairs. This was further confirmed by the estimation of a grand average Event-Related Potential (ERP; see Supplemental Figure S3). Let us note that volume conduction prevents from affirming that the recordings only reflect ventral STN LFP. Consequently, all subsequent interpretations apply to the global STN region.

#### 2.5.2. Time-frequency analyses

Before time-frequency analyses, we removed overall phase-locked activity by subtracting the ERP. Critically, given that the time-frequency analyses we used can produce edge-effects, these were all performed after a data-reflection procedure. In brief, a mirrored version of the signal was concatenated before and after the original signal, resulting in 9 seconds signal (instead of 3 seconds) before time-frequency decomposition. The resulting signals were trimmed back to their original size after the decomposition, thereby avoiding edge-effects on power and intertrial phase clustering estimates.

Complex Morlet wavelet convolution in the frequency domain was applied for time-frequency decomposition. Frequencies from 1 to 40 Hz were studied, while frequencies higher than 40 Hz were not investigated, since cognitive and emotional processing have been so far associated mostly with frequencies up to the beta range. Convolution was performed by multiplying the Fourier transform of the LFP signal by the Fourier transform of a set of complex Morlet wavelet defined as:

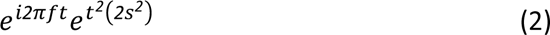

and by taking the inverse fast-Fourier transform of the result. In Equation (2), *t* is time, *f* is frequency ranging from 1 to 40 Hz in 50 logarithmically spaced steps. *S* corresponds to the width of the wavelets and is defined as:

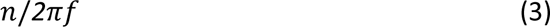

The parameter *n* is the number of cycles, which directly impacts the compromise between frequency and temporal resolution. A greater value of *n* will favor frequency resolution over temporal resolution, while a lower *n* will result in the opposite. We used a logarithmically increasing *n* from 4 to 10, thus favoring temporal resolution for lower frequencies, and frequency resolution for higher frequencies, which is indicated when one investigates condition-specific transient changes (Cohen, 2014).

Spectral power was obtained by taking the squared magnitude of the resulting complex analytic signal (z) at each time point t:

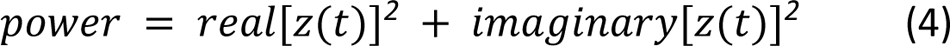

In order to be able to compare conditions, a decibel transform was applied to normalize power results as follows:

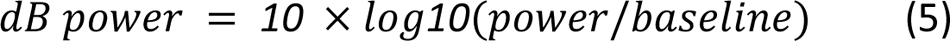

Baseline power was computed from −500 to −200 ms before stimulus onset. Average baseline power was calculated over all trials, irrespective of condition using that time-window.

Synchronization of STN oscillations across trials was also investigated by the means of intertrial phase clustering (ITPC). This measure ranges from 0 to 1, and quantifies the similarity of phase across trials, with 0 indicating a uniform phase angle distribution and 1 indicating perfect phase clustering. ITPC was computed as follows:

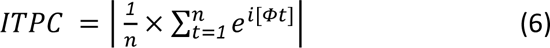

In Equation (6), *n* is the number of trials, and Ф is the phase angle at trial t. Since all conditions did not have the same trial count, ITPC was normalized to ITPCz (in arbitrary units) in order to be able to compare conditions (see Cohen, 2014):

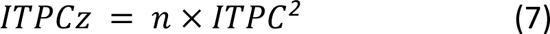

In Equation (7), *n* is the number of trials.

Further analyses were based on power and ITPC extracted from relevant time-frequency windows, for which selection was a two-step process. First, significant changes from baseline were investigated by permutation analyses (Maris and Oostenveld, 2007). For each subject and at each of the 1000 iterations, the time-frequency map was cut at a random point in time and the second half was placed before the first half, thus resulting in a complete misalignment of the time of stimulus onset. This procedure resulted in a distribution of time-frequency maps under the null hypothesis that power/ITPCz was similar across time. A z-score transformation was then applied on the real power/ITPCz data by subtracting the average value under the null hypothesis, and dividing by its standard deviation. Correction for multiple comparisons was carried out using cluster-based correction. The significance threshold was set at p = 0.05.

After permutation testing, smaller time-frequency windows of interest were defined in significant clusters by visual inspection. Critically, we avoided circular inference by performing this selection on grand average maps. Hence, selection was blind to subject- or condition-specific changes in power/ITPCz, and was thus orthogonal to the effects investigated. Averaged power/ITPCz was extracted from these windows for further analyses.

Investigation of condition effects on power and ITPCz were performed using linear mixed models. Model selection was performed in the same way as described in Section 2.4. In addition, compliance with the assumptions of normality and homogeneity of variance of the models’ residuals was checked by visual inspection.

For all power and ITPCz analyses, the following model equation was selected:

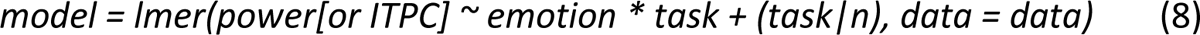

However, for the ITPCz analysis focusing on one of the delta windows, a lower AIC was found when adding the nucleus (left or right) factor (although nucleus did not significantly impact ITPCz after inspection of the results):

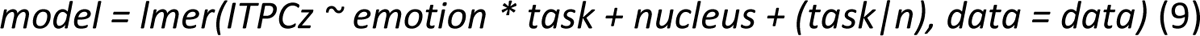

For all models, significance of fixed effects was evaluated by the *anova*{stats} function that provides p-values based on F tests. Marginal and conditional R² were also calculated for all models. When applicable, *post-hoc* tests were carried out using Tukey tests computed by the *glht* function of the {multcomp} package, which computes adjusted p-values using individual *z* tests (Hothorn et al., 2008). Each model significance threshold was adjusted using Bonferroni correction when multiple time-frequency windows were investigated, with p = 0.05/number of time-frequency windows.

### 2.6. LFP-behavior relationships

We investigated whether LFP time-frequency variables (power/ITPCz) were associated with patients’ accuracy at the group level. We used Pearson correlations between accuracy and power/ITPCz extracted from the relevant time-frequency windows. Correlations were also estimated between the difference in power/ITPCz between neutral and fearful expressions and the difference in accuracy between neutral and fearful expressions to investigate emotion effects associations.

## 3. Results

### 3.1. Behavioral results

Figure 2 presents accuracy as a function of stimulus emotional expression and task type. The accuracy rate appeared to be higher for the explicit task (0.91, sd = 0.28) than the implicit task (0.88, sd = 0.32), which was confirmed by a significant task effect (*X^2^* = 7.03, p = 0.008; conditional R^2^ = 0.15, marginal R^2^ = 0.01). Conversely, the emotional stimulus was not associated with changes in accuracy (*X^2^* = 0.3, p = 0.58), nor did it influence the effect of task (*X^2^* = 0, p = 0.99). Overall, accuracy was higher for the explicit than the implicit task.

**Figure 1.**
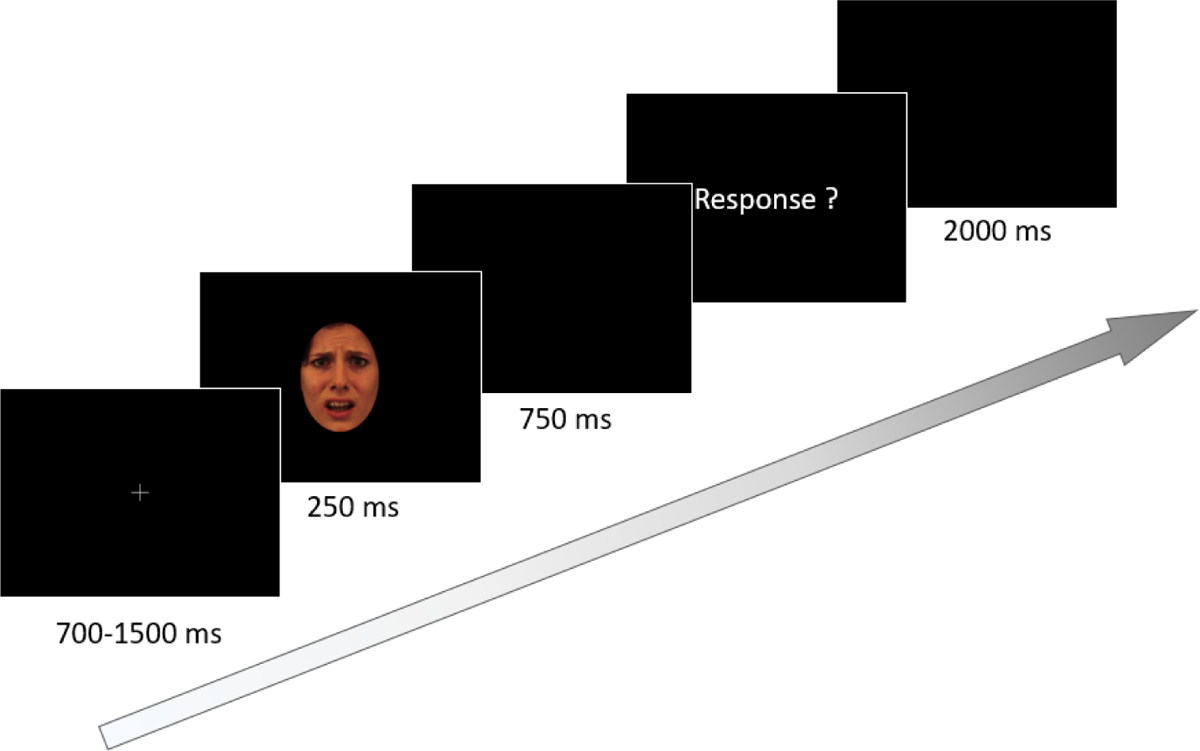
At the beginning of each trial, a central fixation cross was displayed for 700-1500 ms. Then, the stimulus (man/women displaying neutral or fearful expression) appeared for 250 ms. After 750 ms, patients were prompted to respond. After the response, a black screen was displayed for 2000 ms before the start of the following trial.

**Figure 2.**
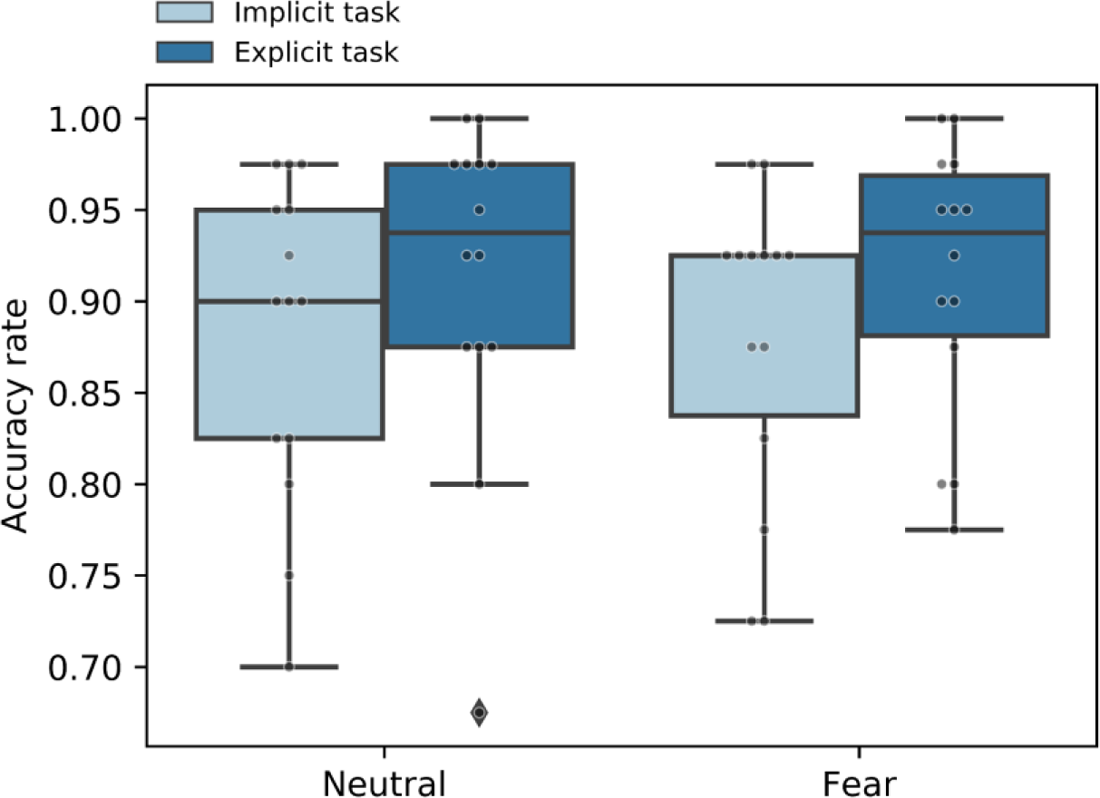
Boxplots of accuracy rate as a function of stimulus emotional valence and task type. Each data point corresponds to the average accuracy of a subject.

### 3.2. Power results

Power analyses focused on 3 different time-frequency windows (Figure 3): (i) A first window in the delta range that showed a significant increase in power compared to baseline whatever the experimental condition (from 2.123 to 3.596 Hz, from 490 to 760 ms after stimulus onset; frequency of peak power: 2.8 ± 0.6 Hz), (ii) a second window in the alpha range with a significant decrease in power (from 8.25 to 12.93 Hz, from 1435 to 1780 ms; frequency of peak power: 10.1 ± 1.9 Hz), and (iii) a third window in the beta range with a significant decrease in power as well (from 15.03 to 27.45 Hz, from 1390 to 1810 ms; frequency of peak power: 21.01 ± 5.1 Hz). Since we investigated three different time-frequency windows, we used a corrected significance threshold of p = 0.05/3 = 0.016. Overall, we found no significant difference in power between the left and right STN (F_(1, 12272)_ = 0.15, p = 0.69).

**Figure 3.**
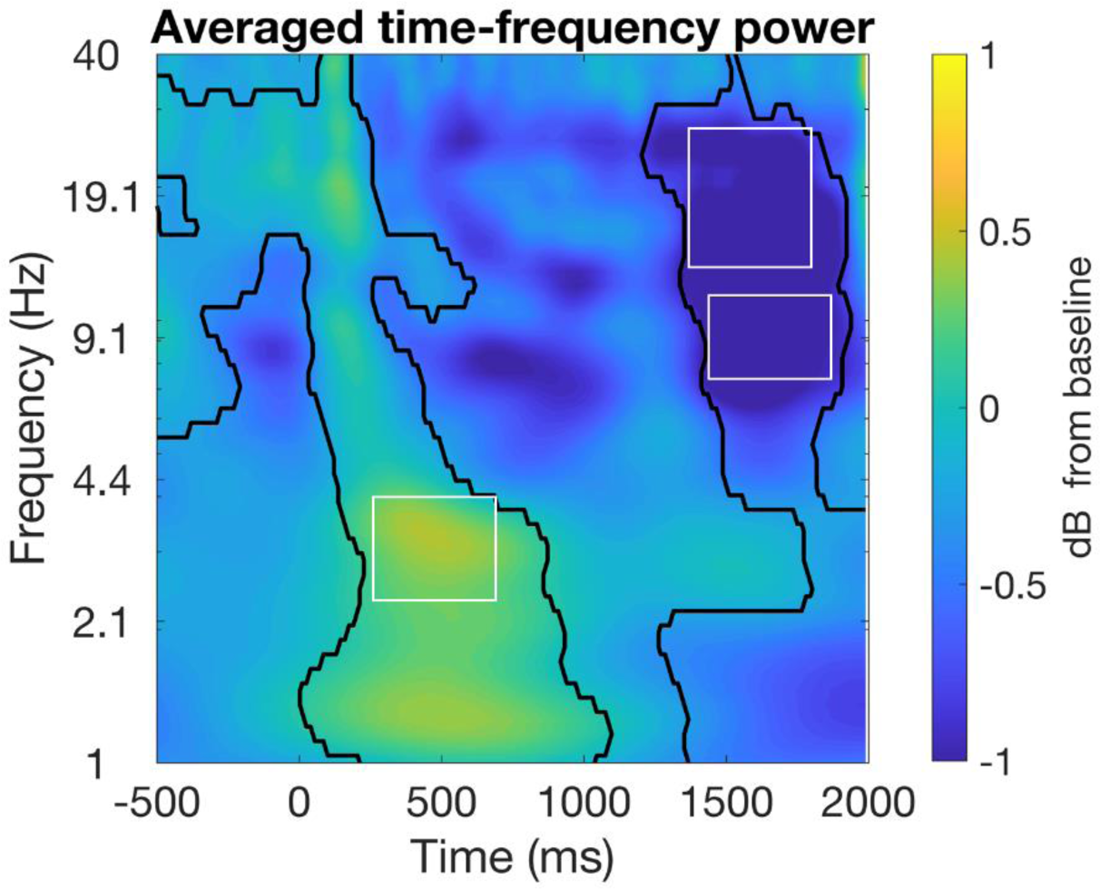
Global average time-frequency power map. Time 0 corresponds to the onset of the stimulus. Black contour lines indicate the regions of significant changes from baseline after permutation analyzes with cluster correction. The plain line rectangles correspond to the time-frequency windows chosen for the analyses: delta, alpha, and beta bands.

#### 3.2.1 Delta window

Figure 4.A illustrates power extracted from the delta time-frequency window. No clear changes in power could be observed according to either emotional expression or to task type (see also Table 2 for all condition-specific results). Statistical analyses revealed no significant effect of emotion (F_(1, 4036)_ = 0.63, p = 0.42; conditional R^2^ = 0.07, marginal R^2^ = 0.006), nor task (F_(1, 27)_ = 2.7, p = 0.11), nor interaction between the two factors (F_(1, 4036)_ = 1.4, p = 0.24). Overall, STN delta power seemed unaffected by changes in emotion or task.

**Figure 4.**
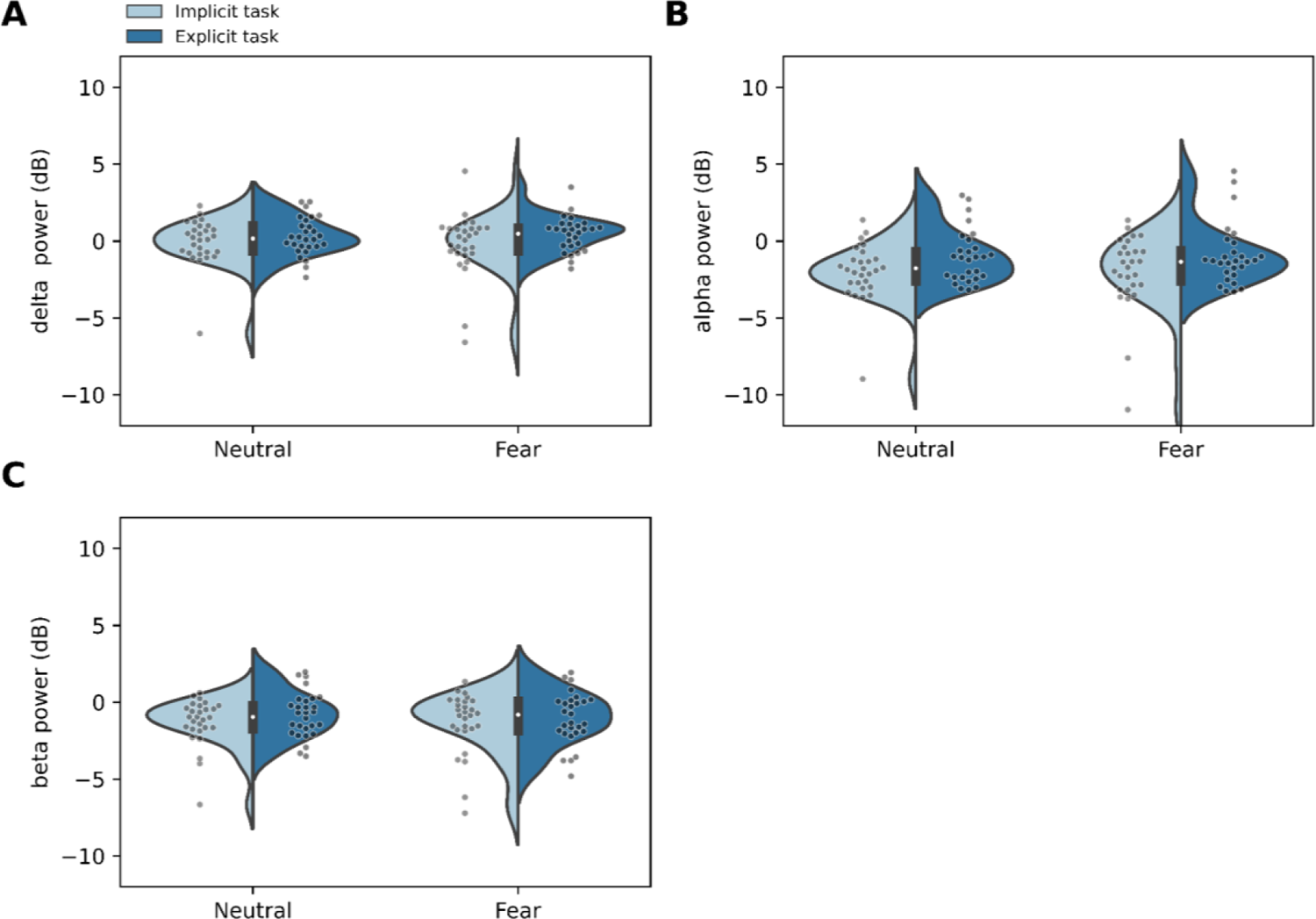
Power as a function of stimulus fearful/neutral emotional expressions and task for delta (A), alpha (B), and beta (C) time-frequency windows. Violin plots present data distribution with an inset boxplot. Each data point corresponds to the average power of a nucleus.

**Figure 5.**
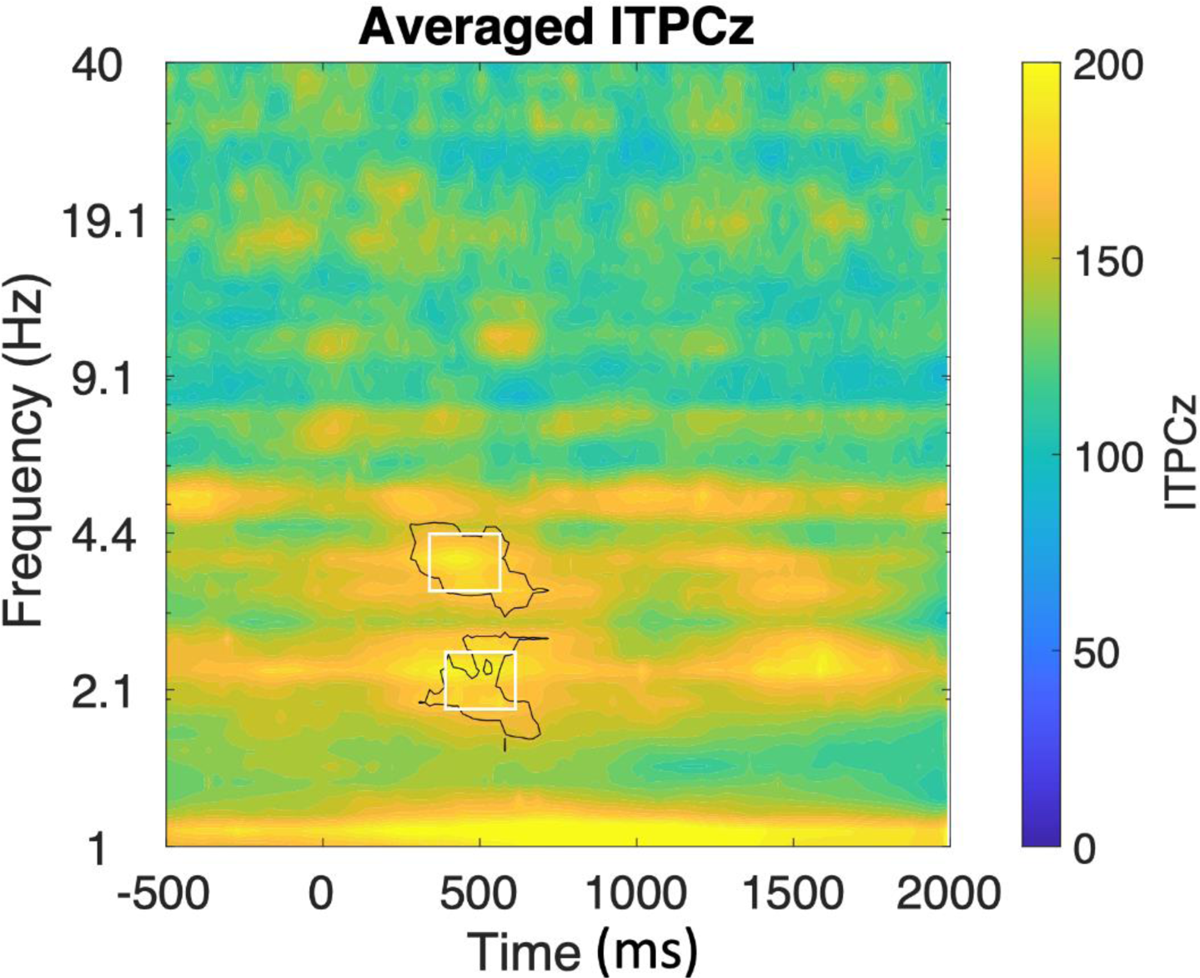
Global average time-frequency ITPCz map. Time 0 corresponds to stimulus onset. Black contour lines indicate the regions of significant changes from baseline after permutation analyzes with cluster correction. The plain line rectangles correspond to the time-frequency windows chosen for the analyses: delta and theta bands.

**Table 2:**
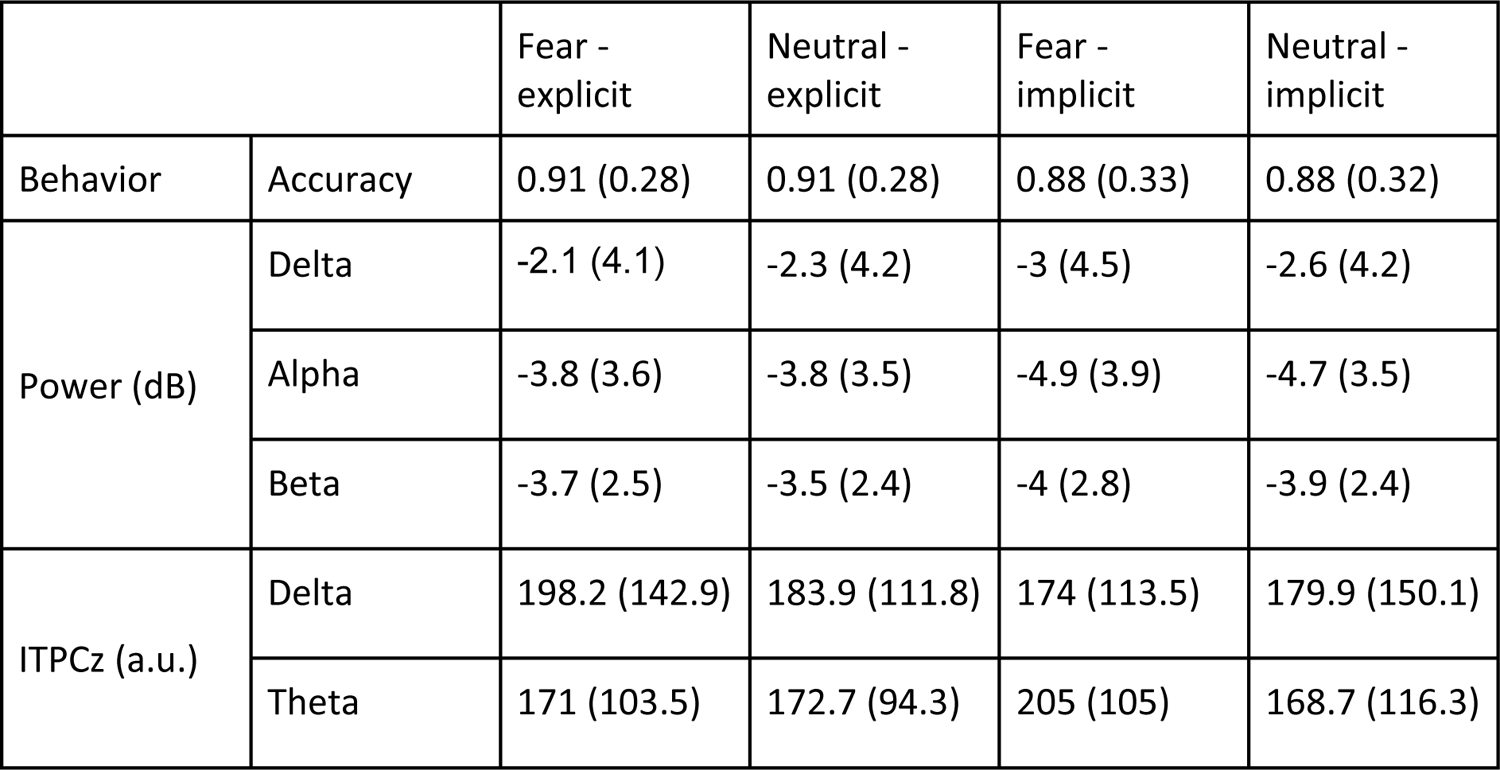
Behavioral and LFP results showing mean (SD) accuracy rate, power and ITPCz extracted in the different time-frequency windows, according to the emotional expression of the stimulus and the type of task.

|  |  | Fear - explicit | Neutral - explicit | Fear - implicit | Neutral - implicit |
| --- | --- | --- | --- | --- | --- |
| Behavior | Accuracy | 0.91 (0.28) | 0.91 (0.28) | 0.88 (0.33) | 0.88 (0.32) |
| Power (dB) | Delta | -2.1 (4.1) | -2.3 (4.2) | -3 (4.5) | -2.6 (4.2) |
|  | Alpha | -3.8 (3.6) | -3.8 (3.5) | -4.9 (3.9) | -4.7 (3.5) |
|  | Beta | -3.7 (2.5) | -3.5 (2.4) | -4 (2.8) | -3.9 (2.4) |
| ITPCz (a.u.) | Delta | 198.2 (142.9) | 183.9 (111.8) | 174 (113.5) | 179.9 (150.1) |
|  | Theta | 171 (103.5) | 172.7 (94.3) | 205 (105) | 168.7 (116.3) |

#### 3.2.2 Alpha window

Similarly to the delta window, power extracted from the alpha time-frequency window was similar, whatever the emotion displayed (neutral or fearful) or task (explicit or implicit; Table 2, Figure 4.B). Indeed, there was no significant difference in power according to emotion (F_(1, 4034)_ = 0.01, p = 0.93; conditional R^2^ = 0.24, marginal R^2^ = 0.02), or task (F_(1, 27)_ = 5.43, p = 0.03). The interaction between the emotion and task factors was also not significant (F_(1, 4034)_ = 1.26, p = 0.26). Taken together, these results suggest that alpha oscillations were not modulated by task or emotion.

#### 3.2.3 Beta window

Figure 4.C presents the power extracted from the beta time-frequency window. No clear changes in power were present according to emotional expression or task in beta power (Table 2), since statistical analyses revealed no significant differences between fear and neutral expressions (F_(1, 4033)_ = 2.7, p = 0.09; conditional R^2^ = 0.4, marginal R^2^ = 0.006), as well as between implicit and explicit tasks (F_(1, 27)_ = 0.95, p = 0.34). No significant interaction was found between the two factors (F_(1, 4033)_ = 1.01, p = 0.31). These results suggest that STN beta power was not modified by emotion or by the implicit/explicit nature of the task.

### 3.3. ITPCz results

ITPCz analyses focused on two different time-frequency windows. The first window was in the delta range (from 1.969 to 2.661 Hz, from 370 to 595 ms after stimulus onset; frequency of peak ITPCz: 2.3 ± 0.3Hz), a second window was in the delta-theta range (from 3.335 to 4.18 Hz, from 300 to 580 ms; frequency of peak ITPCz: 3.8 ± 0.3Hz). To ease the reading, we further refer to this delta-theta window as theta window. Since we investigated two different time-frequency windows, we used a corrected significance threshold of p = 0.05/2 = 0.025.

#### 3.3.1. Delta window

ITPCz extracted from the delta time-frequency window is presented in Figure 6.A. ITPCz was similar between the two emotional stimuli, which was confirmed by the statistical analyses that revealed no significant emotion effect (F_(1, 81)_ = 0.008, p = 0.93; conditional R^2^ = 0.41, marginal R^2^ = 0.005), nor task effect (F_(1, 81)_ = 0.68, p = 0.41). No significant interaction was found between task and emotion (F_(1, 81)_ = 0.32, p = 0.57). Overall, this suggests that delta phase alignment across trials was similar in all experimental conditions.

**Figure 6.**
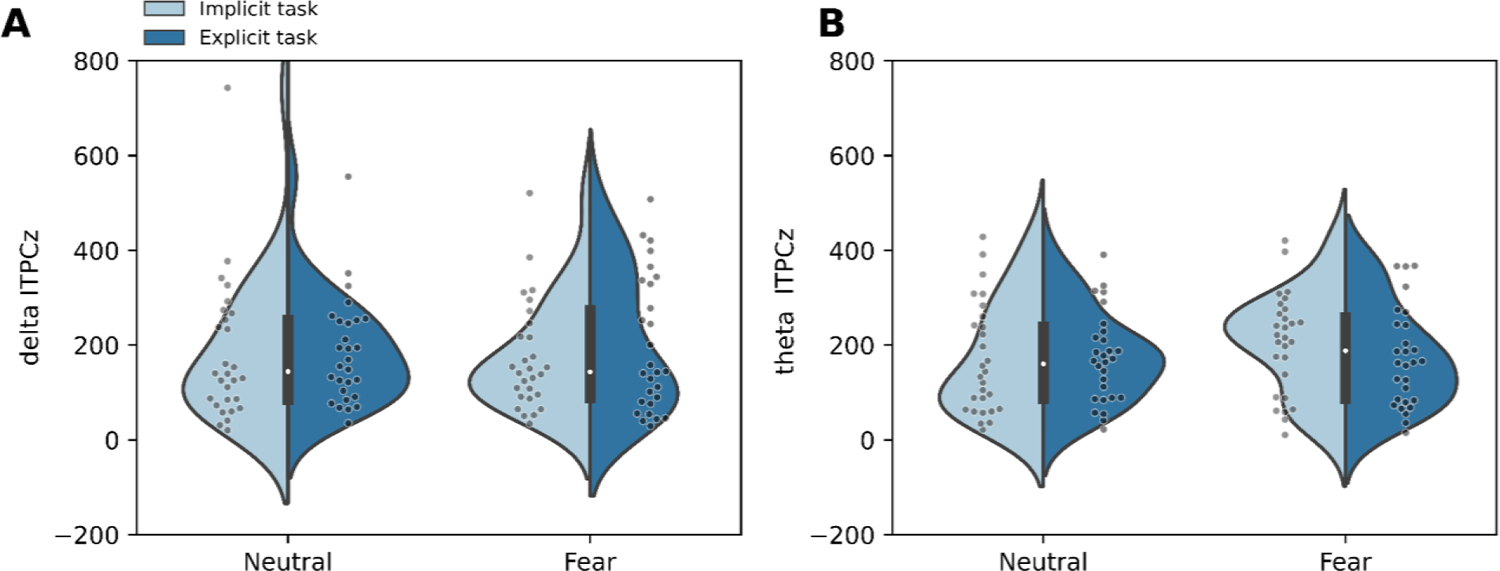
ITPCz as a function of stimulus emotional expression and task type for delta (A), and theta (B) time-frequency windows. Violin plots present data distribution with an inset boxplot. Each data point corresponds to the average ITPCz of a nucleus.

#### 3.3.2. Theta window

A similar pattern of results than for delta was observed for theta ITPCz. Figure 6.B illustrates that theta ITPCz was similar for both the implicit than the explicit task and that no changes according to emotion were observed. This was confirmed by the statistical analyses showing no emotion (F_(1, 81)_ = 1.17, p = 0.28; conditional R^2^ = 0.6, marginal R^2^ = 0.001) or task effects (F_(1, 81)_ = 0.17, p = 0.68). No significant interaction was found between task and emotion effects (F_(1, 81)_ = 2.71, p = 0.1). These results suggest that phase clustering in the theta band was unaffected by the emotion or the task’s nature.

### 3.4. LFP-behavior relationships

Each power/ITPCz time-frequency window was investigated for potential association with overall task accuracy. We also computed correlations between the difference in accuracy according to emotion and the difference in power/ITPCz according to emotion to specifically investigate emotion effects associations. Since power was extracted from three windows, and ITPCz from two windows, the same statistical significance thresholds than previously were applied: p = 0.016 and p = 0.025 for models using power and ITPCz, respectively.

The only significant correlation was found between theta ITPCz and accuracy (Pearson’s r = 0.23, p = 0.016), suggesting that increased theta phase clustering was associated with increased accuracy. All other correlations were not significant (p>0.05). Scatterplots can be found in supplementary figures S4-5.

We also calculated correlations between, accuracy, power and ITPCz data and clinical variables. No significant correlation was found between accuracy and clinical data.

## 4. Discussion

Here, we aimed at characterizing the modulation of neural oscillations in response to emotional faces depending on the orientation of attentional resources using explicit (emotional categorization) and implicit (gender categorization) tasks. Considering the STN central place in motor, cognitive, and limbic functions; we expected changes in STN neural oscillatory dynamics in different frequency bands in response to emotional faces (alpha band) in different attentional orientation situations (theta and beta bands). Finally, we expected that these changes would be associated with behavior. However, our results did not allow us to conclude on the STN involvement in the interaction between emotional and cognitive processing.

### 4.1. Accuracy depends on the task demands

In order to evaluate STN involvement in explicit and implicit emotional processing, we relied on a task that puts the participant in two different attentional situations: they were asked to either discriminate between two facial emotions (explicit task), or to recognize face gender (implicit task). In both situations, facial emotion could vary between neutral faces and fear-expressing faces. Patients in our study were equally accurate whether the faces expressed fear or a neutral expression. However, patients were less accurate when they had to focus on the non-emotional attribute of the stimulus (gender). These results could be interpreted as the consequence of an interfering effect of emotional stimuli on gender categorization, which would be in line with previous observations proposing that the saliency of emotional faces could capture attention (Carretié, 2014; Cohen et al., 2016). However, the absence of an emotional valence effect on behavior in our study strongly mitigates this interpretation. Furthermore, the tasks were always presented in the same order (implicit, then explicit). Although this choice was made to prevent patients from focusing at first on the emotions displayed, it also introduced bias because the same faces were used in both tasks. Thus, increased accuracy in the explicit task might also be related to previous knowledge of the faces. Nonetheless, we cannot rule out the possibility that categorizing gender could have been made more difficult by the presence of varying emotional expressions throughout the task. Another way to interpret these results could be that the gender categorization task was inherently more ambiguous and difficult because of the absence of non-facial features such as hair. These attributes are indeed less relevant for the discrimination of facial expression.

We did not observe lower accuracy for fear compared to neutral expressions in the implicit task, which was reported by (Knyazev et al., 2009). This discrepancy is hard to explain, since studies using such tasks are scarce. One possible explanation is that accuracy in our study was not evaluated in the same way for both explicit and implicit tasks. Indeed, in Knyazev et al. (2009), accuracy in the explicit task was dissociated from gender categorization and participants were asked to judge the friendliness of the face instead of a simple emotion categorization. Another possible explanation is that the emotions displayed were not strong enough in the sense that they did not elicit a sufficient arousal. Overall, our results suggest that behavioral performances during the task were more affected by the nature of the task than by stimuli emotional expression at least for neutral and fearful expressions.

### 4.2. Facial emotional expression was not associated with STN power changes

Enhancement of beta oscillations’ power is mostly associated with motor behavior, and is considered to be a neurophysiologic marker of PD related to rigidity and bradykinesia. Nevertheless, beta oscillations are also involved in orienting attentional resources (Buschman and Miller, 2007) and have been suggested to play a role in active inhibition in cognitive and motor contexts (Zavala et al., 2015). Other frequency bands have been associated with cognitive processing in the STN, such as the theta band that has been repeatedly reported to be involved in conflict processing (Cavanagh et al., 2011; Duprez et al., 2019). Regarding emotional stimuli processing, changes in STN oscillatory power were mostly described in the alpha band (Brücke et al., 2007; Huebl et al., 2014). Changes in alpha band power according to emotional processing has also been described for vocal emotional stimuli (Benis et al., 2020). Consequently, we expected changes in STN power for the alpha band, since our task was designed to induce explicit and implicit facial emotional processing by the STN. Our results did not identify emotion or task-related differences in either delta, alpha and beta frequency bands. Indeed, although we did find delta power increase around 500 ms after stimulus onset, and alpha and beta power decrease around 1700 ms after stimulus onset, none of these were modulated by emotion or attentional demand. This suggests that these oscillatory changes compared to baseline were likely linked to overall task execution mechanisms (e.g. movement-related) and non-specific to emotional recognition processes. Furthermore, we did not identify any relationship between STN power at the investigated frequencies and patients’ behavioral performances (on overall accuracy, and on the emotion effect on accuracy). On the one hand, the absence of alpha and beta power modulations by emotional expression or attention focus is surprising, since changes in alpha power have been associated with the processing of pleasant, as compared to unpleasant stimuli. For instance, Huebl et al. (2014) studied STN LFP in 12 PD patients and found that alpha power significantly decreased when patients saw positive or negative arousing stimuli. Differences in power changes between pleasant and unpleasant stimuli were correlated with depressive symptoms in patients. However, it is worth noting that Mandali et al. (2021) only found such alpha changes when focusing on the left STN only. In our results, we didn’t find differences in power between both nuclei. One possible interpretation of the discrepancy between our results and the literature is the presence of differences in task design. For instance, we chose to focus only on one negatively valanced emotion (fear expression) and neutral stimuli. Furthermore, our task involved an active discrimination behavior (emotional expression or gender), while previous STN-LFP studies reporting alpha changes used passive viewing paradigms (Brücke et al., 2007; Huebl et al., 2014; Kühn et al., 2005), with stimuli remaining on the screen for 1-2 s. Hence, our task involved further cognitive processing and motor behavior, which have been related to beta suppression. Of course, another possible explanation might lie in the limited sample size which is applicable to our study and previous STN-LFP research (see limitations section). On the other hand, this implicit/explicit task, which necessarily involves a complex interaction between emotional processing, orientation of attention - cognitive control, has never been performed before with STN-LFP recordings (although it has been with EEG (Yu et al., 2014)). Consequently, the interaction of emotion-, attention-cognitive control-specific STN oscillatory changes would hardly be seen as a simple combination of all previously known dynamics.

### 4.3. STN functional organization did not change with emotional fearful expression and attentional focus

Most studies investigating STN-LFP neural oscillations focused on power results extracted from time-frequency decomposition methods. Aside from a few exceptions (Duprez et al., 2019; Zavala et al., 2016, 2013), the phase information of STN oscillations is either ignored or not reported. However, neural oscillatory phases carry valuable information, especially when studying how phase is consistent across trials of an experiment. Here, we investigated intertrial phase clustering (ITPC), which can be defined as the strength with which the timing of frequency-specific oscillations clusters over trials. This can be interpreted as an index of functional organization, since strong clustering indicates that oscillations have a similar timing across trials (Cohen, 2014). Our results showed a transient increase in functional organization occurring around 500 ms after stimulus presentation in the delta and theta bands, indicating that organization of delta and theta oscillations became temporarily consistent across trials. However, the valence of facial emotion and whether the task was explicit or implicit did not modify neither delta nor theta functional organization. This suggests that functional organization was not modulated by emotion or by the focus of attention. However, further investigations are needed to firmly rule out emotion or task effect on STN functional organization. It is worth noting that the time-frequency window with increased delta ITPC coincides with the window for power increase that we described in delta. Finally, we found evidence that theta functional organization was linked with behavioral performance, since an increase in theta phase clustering was associated with greater tasks’ accuracy. However, this correlation between STN activity and behavior was the only one that was significant and should thus be interpreted with caution.

Although our limited sample size prevents from firm conclusions, our results, suggest that delta and theta functional organization in the STN were not involved in emotion- or attention-related mechanisms during the task. Rather, as this was the case for power, this points to a general task-execution mechanism. Interestingly, delta oscillation modulations in the STN have been reported in situations of cognitive control (Zavala et al., 2013). Delta oscillations have also been associated with cognitive control at the cortical level in EEG studies, especially during errors (Cohen and van Gaal, 2014), or while detecting relevant stimuli when distractors are present (Herrmann et al., 2016). Although it is hazardous to directly compare cortical and STN delta oscillations, the post-stimulus increase in delta and theta ITPC that we observed could be associated with task demands in cognitive control. For instance, although this is purely speculative at this stage, it is possible that the absence of non-facial features made the task more difficult which could have modulated delta/theta phase clustering.

### 4.4. Limitations

The first and most important limitation to this study is related to task design and the absence of emotional-valence effect. Indeed, only one negative emotional valence was used in this study. We chose fear because it is the least well recognized emotion in PD, and was thus well-suited to investigate facial emotional processing in PD (Argaud et al., 2018). Thus, we cannot extend the discussion further on the effect of emotional valence that was found in others studies using positive and negative valences. However, more importantly, the fact that accuracy was not affected by emotional valence suggests that the task was not very sensitive to emotional processing, which prevents strong conclusions on any associated STN activity. Moreover, the decrease in accuracy observed in the implicit task might result from the fact that the implicit task was always presented first (to avoid the focus on emotional features) and the same faces were used in both tasks. Thus, the fact that the faces were familiar in the emotional task might have affected accuracy. Further studying implicit and explicit emotional processing using such task might benefit from balancing conditions and presenting different faces in the task, as well as adding a positively valanced emotion to get a wider view of how power in emotional context varies from a neutral one. Another way to improve this task could be to control for the arousal elicited by the stimuli, which could be done by ratings at the end of the experimental procedure. Also, there was no way to guarantee that the participants’ attention was not towards emotion in the implicit task. This could be important to control but would require controlling for both overt and covert attention. Although overt attention could be controlled by recording saccade and gaze, covert attention to facial features when fixating on other features would be difficult. Indeed, covert attention can be assessed by using flickering elements in the stimulus that would entrain occipital oscillations which could be recorded by EEG (Mora-Cortes et al., 2018). As a whole this would result in a very heavy experimental setup.

Another limitation to our study is the small sample of patients (n = 14). Small samples are however inherent to human LFP studies, given the recruitment difficulties associated with the low number of patients that can undergo STN DBS. Nevertheless, larger samples would lead to more robust and generalizable findings. Furthermore, controlling for medication is very hard in such studies given that recording STN LFP without dopaminergic medication would be very difficult. Nonetheless, we didn’t find any correlation between levodopa equivalent daily dose and STN power or ITPCz. A final issue concerns the association between STN activity and behavior. Indeed, due to technical difficulties, the signal had no response-specific triggers, which prevented an evaluation of the effect of power on accuracy at the trial-level, thereby diminishing statistical power. Following this, the absence of response-specific triggers complicates interpretation of the power decreases we observed, which are likely associated with movement.

## 5. Conclusion

A growing body of literature confirms that the STN is not exclusively a motor territory, but has also an integrative role for cognitive and limbic processes which constantly interact to give rise to the adapted behavior. Our study did not allow us to conclude on the hypothesis that the STN is involved in facial emotional processing, even when emotional information is not relevant to the situation, and attention is directed toward non-emotional stimuli characteristics. Our study suggests that the neuronal oscillatory activity previously described as involved in emotional processing and cognitive control was not affected by emotional fearful expression compared to neutral ones or attentional demand. However, given the low sample size of our study, and the fact that the task did not seem to provide a strong grasp on emotional processing, further investigations are needed to precisely investigate these hypotheses.

## Acknowledgements

The authors would like to thank all the patients who took part in this study. We are also grateful to Christophe Mermoud for his help in setting up the acquisition system.

## Abbreviations

DBS: deep brain stimulation

ITPC: intertrial phase clustering

LFP: Local Field Potential

PD: Parkinson’s disease

STN: subthalamic nucleus

UPDRS: Unified Parkinson’s Disease Rating Scale

S&E: Schwab & England

H&Y: Hoehn & Yahr

LEDD: levodopa-equivalent daily dose

MDRS: Mattis Dementia Rating Scale.

## 6. Supplemental materials

**S1.**
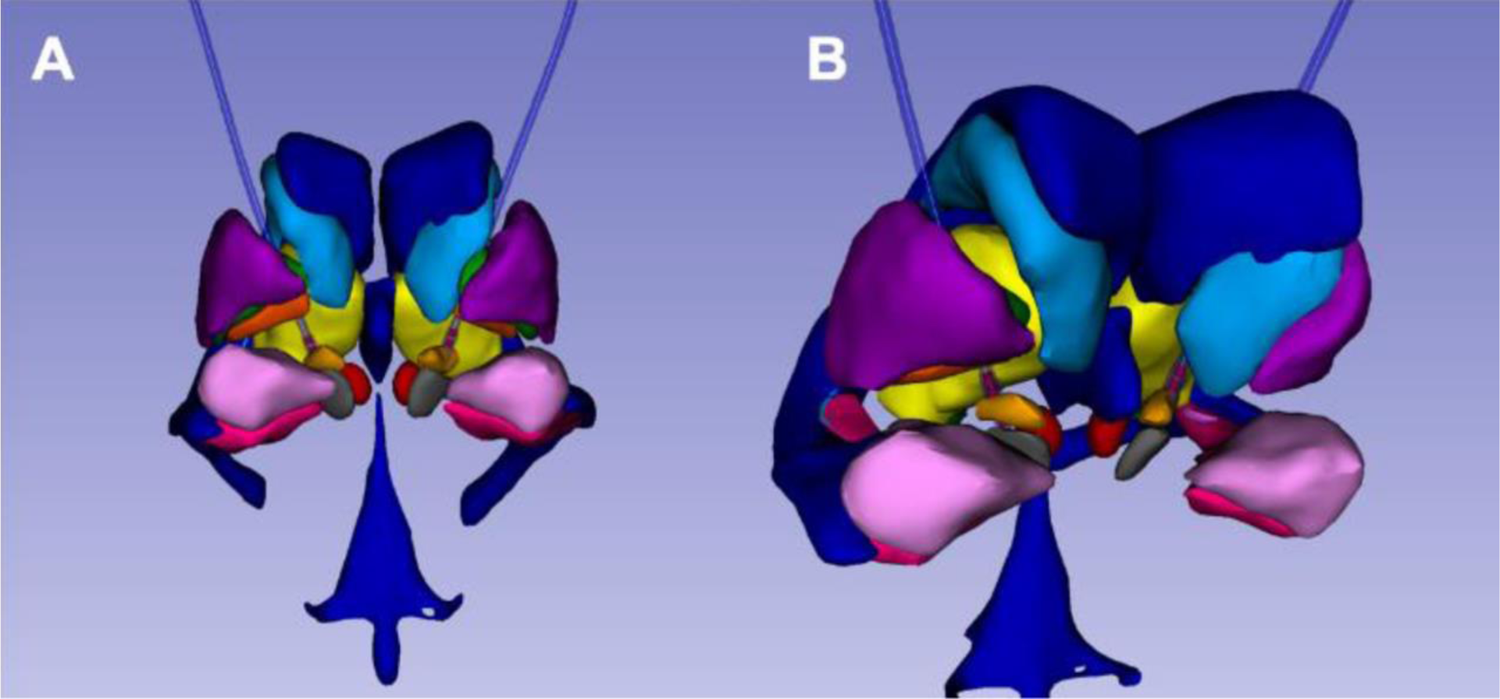
Atlas-based reconstruction showing electrode and contacts position in one patient with the more distal contacts located in the STN (in orange).

**S2.**
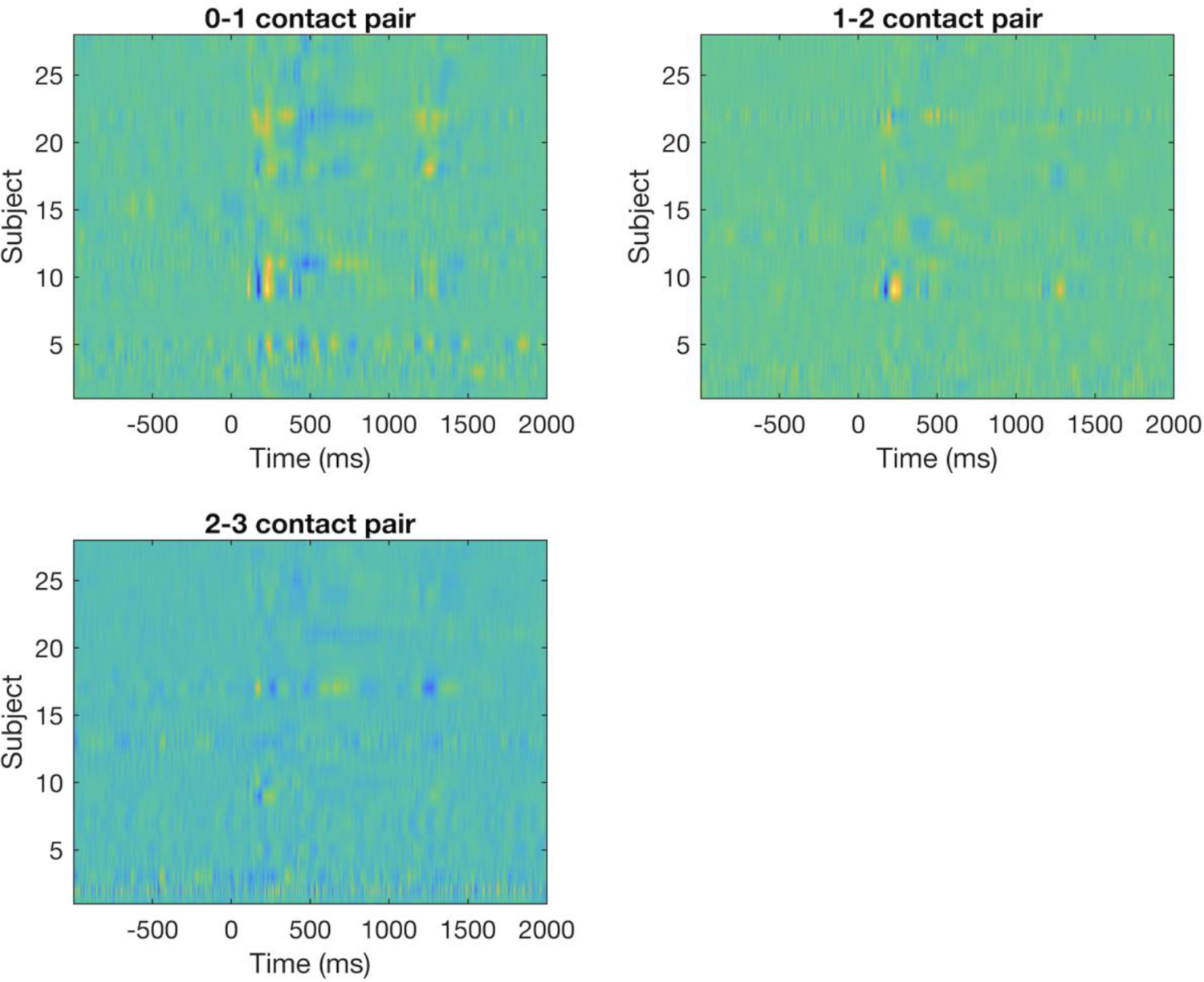
Group-level ERP breakdown of intracranial recordings (ERP computed on all trials) in distal (0-1), intermediate (1-2) and proximal (2-3) pairs of contacts. The marked activity in the distal pair decreased in the intermediate pair of contacts and was merely visible on the proximal pair of contacts. Time 0 represents stimulus onset.

**S3.**
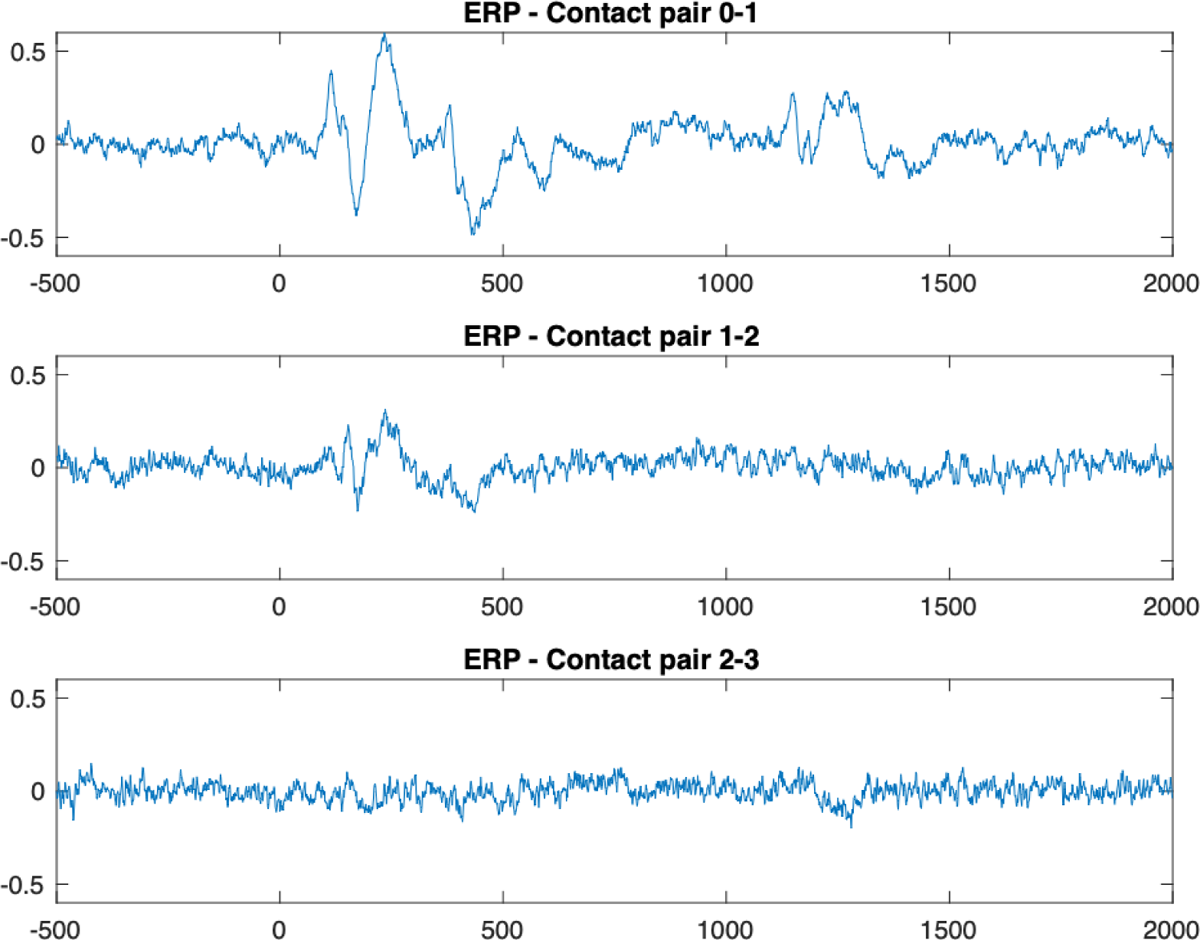
Grand-average ERP breakdown for the three pairs of contacts. A clear ERP can be observed in the 0-1 contact pair, which progressively disappears in other contact pairs.

**S4.**
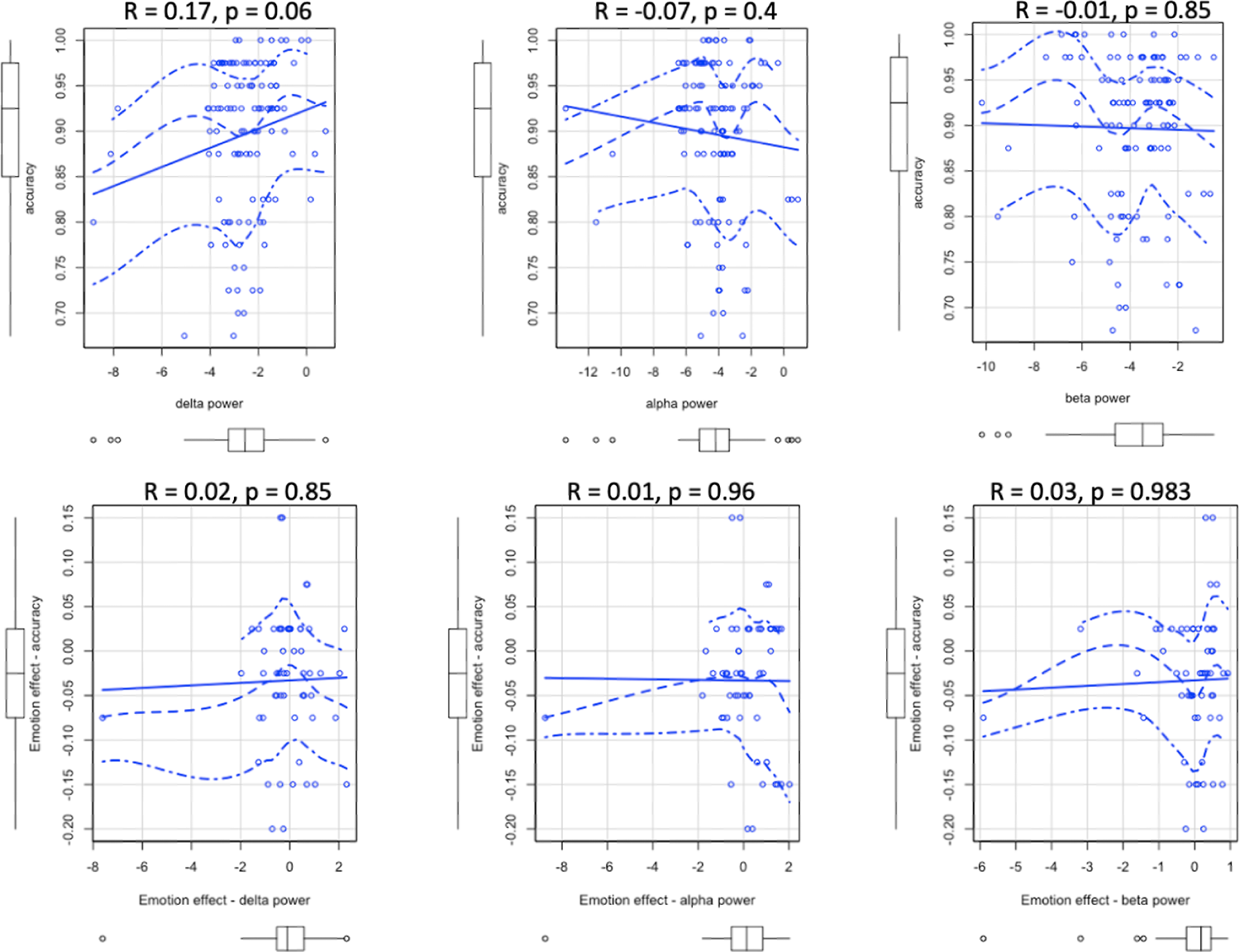
Scatterplots of the correlations between STN power and behavioral measures. Each dot corresponds to a nucleus.

**S5.**
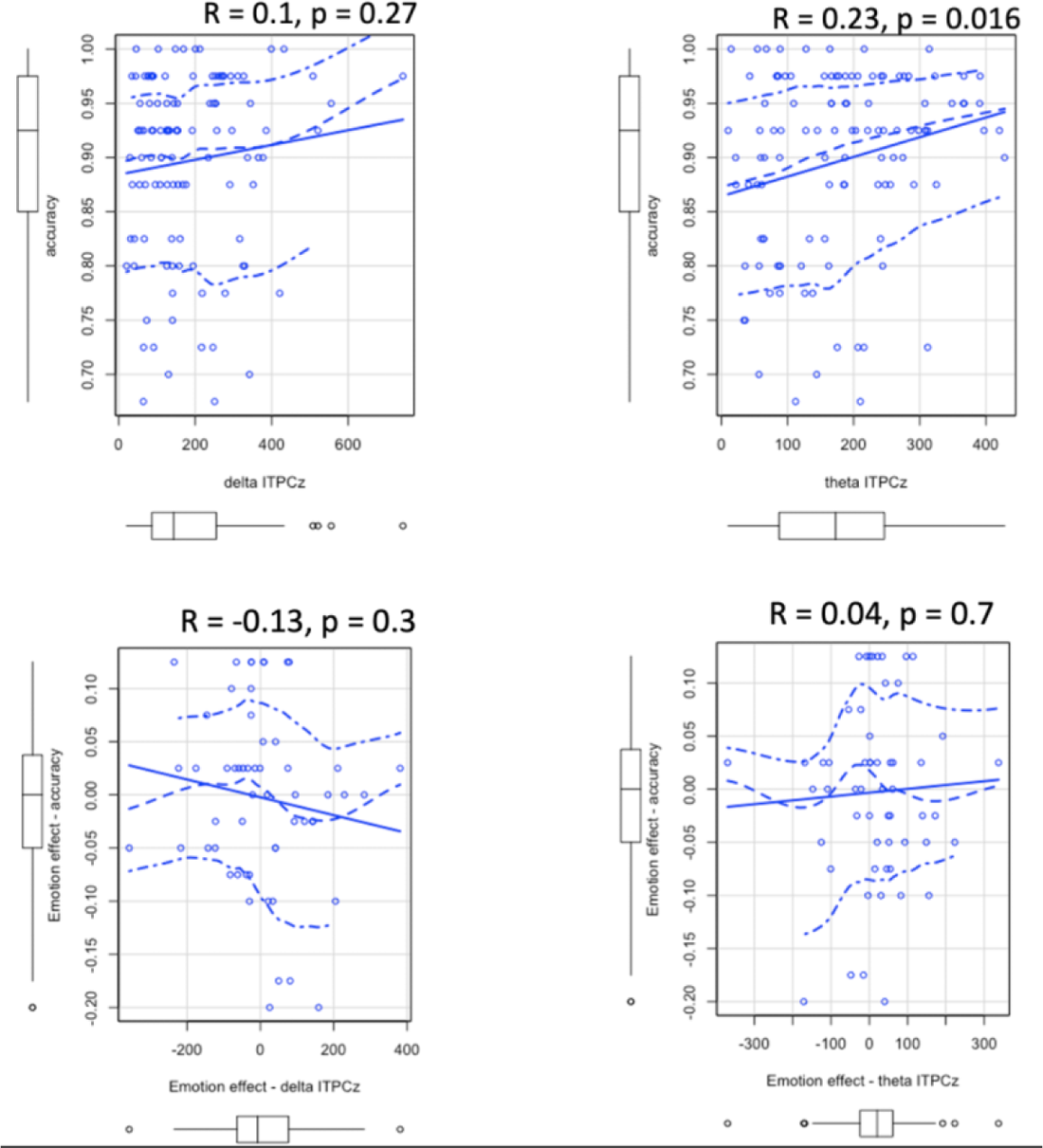
Scatterplots of the correlations between STN intertrial phase consistency and behavioral measures. Each dot corresponds to a nucleus.

